# Characterization and super-resolution imaging of small tau aggregates in human samples

**DOI:** 10.1101/2023.06.12.544575

**Authors:** Dorothea Böken, Dezerae Cox, Melanie Burke, Jeff Y. L. Lam, Taxiarchis Katsinelos, John S. H. Danial, William A. McEwan, James B. Rowe, David Klenerman

## Abstract

Hyperphosphorylation and aggregation of the microtubule binding protein tau plays a key role in the development of Alzheimer’s disease. While the structure of the filamentous aggregates formed in humans has recently been determined to atomic resolution, there is far less information available about the smaller aggregate precursors, thought to be the most neurotoxic. To address this gap, we have developed a single molecule pull-down (SiMPull) able to detect tau aggregates in clinically relevant human samples. This method enables the detection and characterisation of individual tau aggregates, as opposed to averaged features obtained from traditional bulk techniques. We report the number, size and shape of individual aggregates measured via super-resolution microscopy, revealing disease-specific differences in tau aggregate morphology. By adapting the assay to simultaneously detect multiple phosphorylation sites in individual aggregates, we were also able to derive compositional profiles for pathological modifications present in individual aggregates. We demonstrate that tau aggregates in Alzheimer’s disease are significantly more likely to contain both the AT8 and T181 pathological phosphorylation markers, rather than only one. Together, tau SiMPull identified distinct subpopulations of large, modified tau aggregates that were invisible to traditional methodologies. These morphological and compositional differences distinguish samples taken from disease cohorts, offering to illuminate underlying disease mechanisms, and providing a foundation for novel diagnostic strategies.

## Introduction

Abnormal aggregates of tau are a defining characteristic of a range of neurodegenerative diseases collectively known as tauopathies, including Alzheimer’s disease (AD), progressive supranuclear palsy and approximately half of the frontotemporal dementias, including Pick’s disease^1–4^. Tau is an essential protein for the formation and stabilization of microtubules^5,6^. However, hyperphosphorylation of tau causes it to dissociate from microtubules leading to its aggregation^7^. The exact cause of onset remains unknown, however, the formation of tau aggregates begins decades prior to symptoms of cognitive decline. Large insoluble deposits in the form of neurofibrillary tangles (NFTs) containing tau filaments, the product of aggregation, are a primary histopathological hallmark of tauopathies. Indeed, in Alzheimer’s disease, cognitive decline correlates closer with the progression of tau aggregate pathology than with the presence of amyloid-β plaques^8–10^. However, small soluble aggregates are thought to exert potent cellular toxicity^11,12^ and appear to spread from cell to cell as a potential mechanism for the propagation of tau pathology throughout the brain^13,14^.

The structure of the ordered part of tau filaments formed in several tauopathies has recently been determined using Cryo-EM^15,16^. Furthermore, very sensitive tau seeding assays have been developed to detect the number of seed competent tau aggregates in post mortem brain samples or CSF^17–19^. However, despite their importance, the smaller tau aggregates that form during the aggregation process are more challenging to study since they are highly heterogenous in size, shape, and phosphorylation state^20^.

Here, we present a method to study tau aggregates using an adaptation of the single-molecule pull-down (SiMPull) method which combines antibody-based immunoprecipitation with single-molecule fluorescence imaging^21^. We show that this assay quantifies tau aggregates with high specificity and sensitivity in disease-derived samples, including brain tissue homogenates and serum extracts. We report the ability of SiMPull to discern between tauopathy disease-derived and age-matched control samples based on aggregate number. Finally, using co-localization and super-resolution microscopy to characterize aggregate size, shape, and composition we identify disease-associated aggregate subpopulations. Together, these characteristics confirm tau SiMPull as a less-invasive diagnostic tool which can distinguish aggregate subpopulations to track disease pathology and progression in diverse biofluids.

## Results

### Establishing a single-molecule pull-down (SiMPull) for tau aggregates

The single-molecule pull-down (SiMPull) assay is an antibody-based technique that uses fluorescence microscopy to detect and characterize single particles. Antibodies immobilized on a PEG-passivated surface are used to capture tau, which subsequently can be detected using a fluorescently labelled antibody and TIRF microscopy (Fig. 1A). For the specific detection of tau aggregates, SiMPull assays were developed using matched monoclonal tau antibodies for both capture and detection. We reasoned that this configuration required the presence of two identical epitopes within a single particle, as capturing the particle would occupy one binding site and require a second site for binding of the detection antibody. This ensures that the detected tau species are, at a minimum, dimeric. Further, it allows multimeric tau to be isolated without relying on conformation specific antibodies, thereby remaining agnostic to the structure of the tau aggregates. We initially selected two tau antibodies that are commonly used in the field: the phospho-tau specific antibody AT8 (p-Ser202, p-Thr205) and the total tau antibody HT7^22^.

**Fig. 1.**
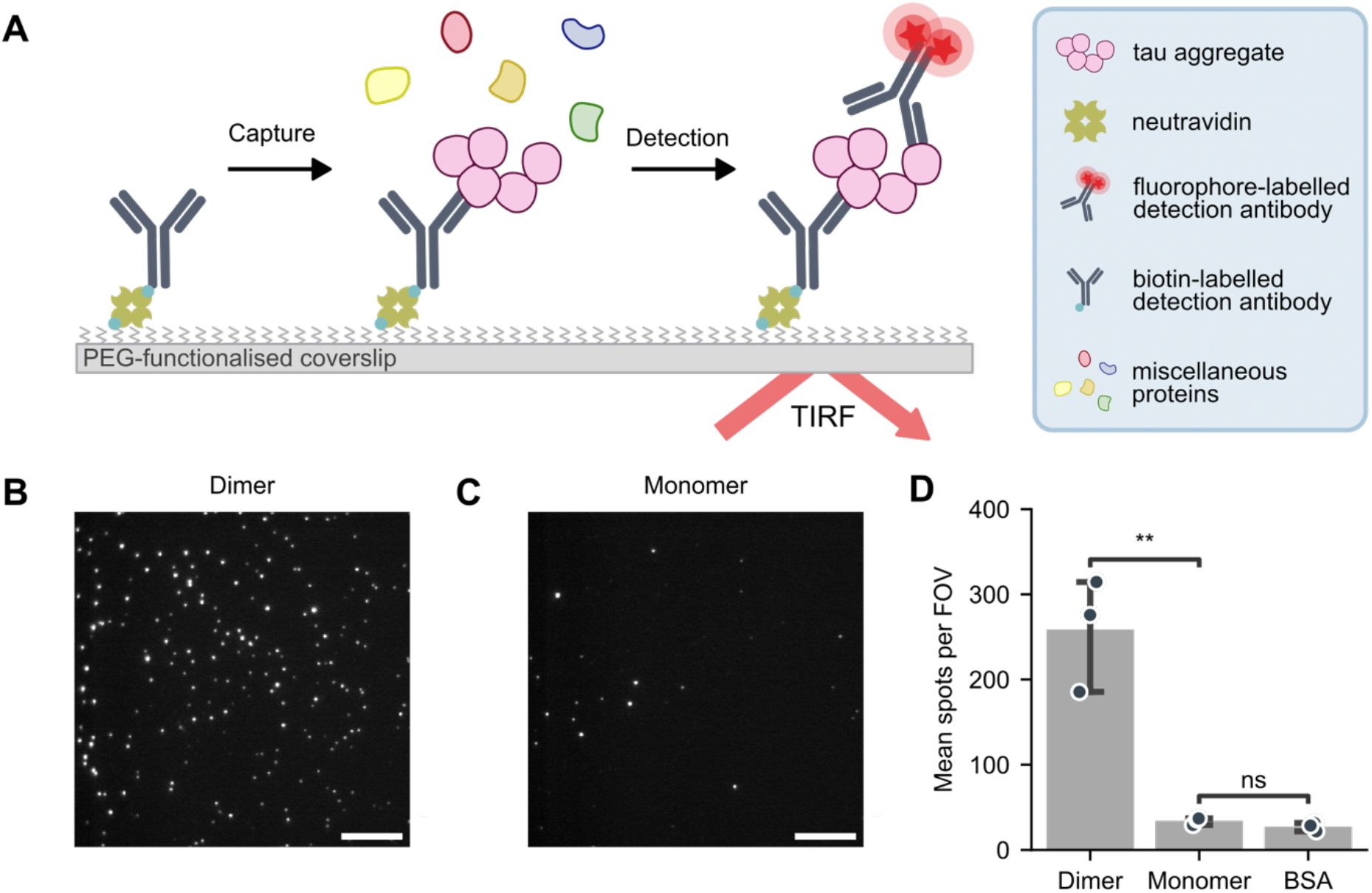
Tau single-molecule pull down (SiMPull) aggregate assay specifically detects multimeric particles. **(A)** Schematic representation of the single-molecule pull down (SiMPull) assay. For the detection of aggregates the same antibody is used for capture and detection. Images are acquired using total internal reflection fluorescence (TIRF) microscopy. **(B)** Representative image of the tau SiMPull assay applied to a dimer-mimicking peptide, containing two linked HT7 epitope sequences. Scale bar = 10 μm. **(C)** Representative image for SiMPull of a monomer-mimicking peptide, containing one HT7 epitope and a randomized sequence. **(D)** Diffraction-limited quantification of the number of spots in individual fields of view. Panel D shows mean ± S.D. of n = 3 technical replicates, compared using a one-way Anova with post-hoc Tukey HSD test. ns: *p* > 0.05, **: *p* < 0.01.

We first sought to confirm the specificity of the assay for tau multimers over monomers. For this purpose, we created synthetic peptides comprising two 10-residue sequences joined via a PEG linker. One peptide contained a single copy of the HT7 epitope tethered to a randomized version of the epitope sequence; this is designated as the ‘monomeric’ peptide given it mimics the presence of the single HT7 binding site available in monomeric tau protein. The second peptide, designated the ‘dimeric’ peptide, contained two HT7 epitope sequences tethered together, mimicking the presence of two HT7 binding sites in multimeric tau species. Importantly, these peptides maintain comparable physicochemical properties to one another owing to their overall identical composition. As a negative control, we used bovine serum albumin (BSA), a well-characterized recombinant control protein which does not contain any HT7 binding sites. These samples were compared via the SiMPull assay, where the primary assay readout was the number of fluorescent spots visible in diffraction-limited images (Fig. 1B, C and Supplementary Fig. 1A).

As expected, the number of fluorescent spots was dependent on the presence and concentration of the HT7 epitope (Fig. 1B and C). Even at very high sample concentrations (0.2 mg/mL), we detected significantly fewer spots in the presence of the monomer-mimicking peptide (34 ± 4 spots) per field of view (FOV) compared to the dimer mimic (260 ± 66 spots per FOV; Fig. 1D; one-way Anova with post-hoc Tukey HSD test, *p* = 0.0084). In contrast, there was no significant difference in the number of spots between the monomer-mimicking peptide and the BSA negative control (*p* = 0.29). The HT7 dimer-mimicking peptide was not detected by an antibody targeting a different tau epitope (AT8; Supplementary Fig. 1B). These data confirmed the assay is able to distinguish tau multimers over monomers.

We further confirmed the specificity of the assay by comparing the detection of (phosphorylated) tau aggregates against other relevant recombinant protein aggregates, namely amyloid-β and α-synuclein aggregates produced *in vitro* (Supplementary Fig. 1C and D). We found on average 12 ± 4 spots per FOV when performing the tau SiMPull assay on these control samples, which is equivalent to the buffer control without any protein present. There was no significant difference between PBS and amyloid-β or α-synuclein (AT8 assay: one-way ANOVA *p* = 0.000092; post-hoc Tukey HSD test, *p* = 1.0 for PBS vs amyloid-β, *p* = 1.0 for PBS vs α-synuclein; HT7 assay: one-way ANOVA *p* = 0.0000050; post-hoc Tukey HSD test, *p* = 0.98 for PBS vs amyloid-β, *p* = 1.0 for PBS vs α-synuclein). Further, we ensured that the recombinant aggregates can be detected with their appropriate antibody combination using corresponding SiMPull assays (Supplementary Fig. 1E and F). Taken together, these results confirmed that tau SiMPull assays can be used to specifically detect tau multimers.

### Diffraction-limited SiMPull detects disease associated differences in tau aggregates

We proceeded to test a complex, biologically relevant protein mixture where we expect a high concentration of tau aggregates, namely homogenized human brain tissue from donors with clinical and neuropathological evidence of the Alzheimer tauopathy and control donors without Alzheimer’s disease. We confirmed the samples contain tau (not aggregate specific) using a commercial enzyme-linked immunosorbent assay (ELISA) (Supplementary Fig. 1G) prior to quantifying their tau aggregate content using the HT7 and AT8 tau SiMPull assays (Fig. 2A and B). On average we detected 600 ± 150 HT7-positive spots and 320 ± 110 AT8-positive spots per FOV (Fig. 2C). We used the concentration of total tau to determine the sensitivity of the SiMPull assays; the limit of detection for the HT7 and AT8 assays was calculated as 874 pg/mL and 2201 pg/ml of total (monomeric) tau respectively ^23^ (Supplementary Fig. 1H). Importantly, we anticipate aggregated tau to comprise only a fraction of the total tau present, and not all tau to be AT8 phosphorylated, meaning the limit of detection for tau aggregates can be expected to be several folds lower in both assay configurations.

**Fig. 2.**
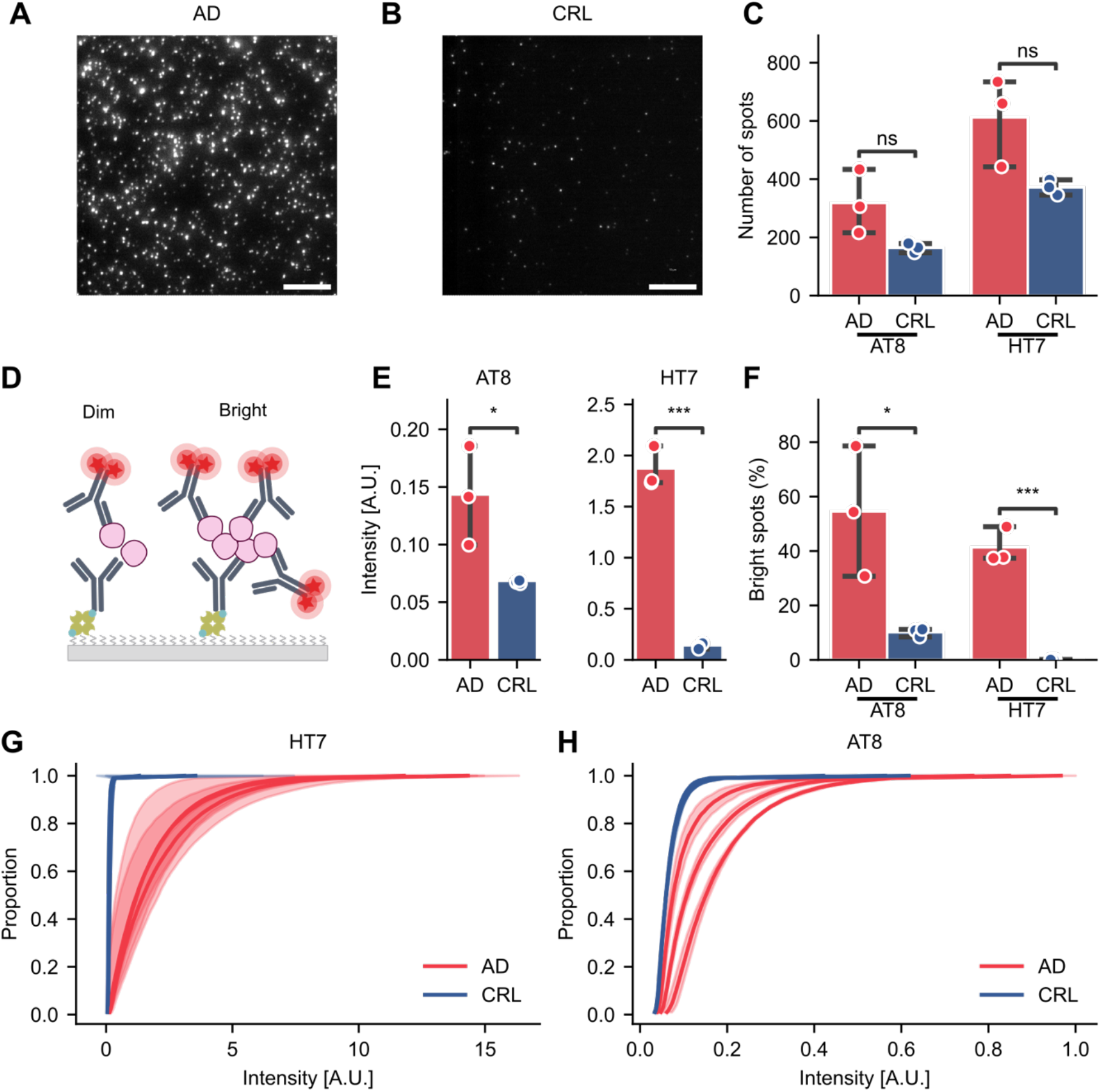
Quantification and characterization of tau aggregates in human brain tissue. **(A)** Representative image obtained for an AD-derived sample using AT8. Scale bar = 10 μm. **(B)** Representative image obtained from a control sample using AT8. **(C)** Quantification of total tau (HT7) and p-tau (AT8) aggregates from AD (Braak stage VI, n = 3) and age-matched control patients (n = 3). **(D)** Schematic of detection antibody binding correlated with aggregate size. Mean intensity of aggregates relates to the number of detection antibodies bound and thereby the aggregate size. **(E)** Mean intensity of tau aggregates in AD, CRL brain and BSA. **(F)** Percentage of very bright aggregates (AT8: intensity > 0.1 A.U., HT7: intensity > 1.5 A.U,). **(G-H)** Cumulative distribution of the aggregate brightness using **(G)** HT7 or **(H)** AT8. Panel C, E, F show the mean ± S.D. of n = 3 biological replicates and asterisks refer to t-tests: ns: *p* > 0.05, *: *p* < 0.05, **: *p* < 0.01, ***: *p* < 0.001.

An advantage of the single-particle nature of SiMPull assays is their ability to derive parameters of each particle individually, extending beyond the mere quantification of tau positive particles. This allows for the identification of other potential disease-associated characteristics. When comparing AD-derived brain homogenate (frontal cortex, Braak stage VI) with that from age-matched control donors (frontal cortex, Braak stage 0/I), despite observing on average 1.5x as many tau-positive particles in AD extracts (Fig. 2C), high variability across AD donors resulted in no detectable statistical difference (t-test, HT7: *p* = 0.054, AT8: *p* = 0.072). Interestingly, we observed positive signal in the age-matched control brains also, indicating the presence of small tau aggregates even at Braak stage 0/1 (when there are no tangles present in the frontal cortex).

When imaged under diffraction-limited conditions SiMPull encodes additional information in the mean brightness of each spot. As larger aggregates present more epitopes to which fluorescently labelled antibodies can bind, we reasoned brightness should be correlated to aggregate size (Fig. 2D). The mean brightness of tau aggregates detected in each sample using HT7 or AT8 (Fig. 2E) revealed significant differences between the disease and control cohorts (t-test, HT7: *p* = 0.00012, AT8: *p* = 0.039). A significantly higher percentage of bright spots (AT8: >0.1 A.U., HT7: > 1.5 A.U.) was also observed in the AD samples compared to the control cohort for both SiMPull configurations (Fig. 2F, t-test, HT7: *p* = 0.00041, AT8: *p* = 0.033). This is similarly evident in the cumulative distribution of spot brightness, a comparison that is made possible only by the single-molecule nature of SiMPull. The distribution of spot brightness’ was highly skewed toward dim (∼small) aggregates in control samples, a feature which was consistent across different control donors in both assays (Fig. 2G and H). In contrast, AD aggregate profiles were more variable between donors, though consistently showed a greater range in brightness of the aggregates, suggesting a greater diversity of aggregate sizes is present in these samples (Fig. 2G and H). Overall, these data support the ability of SiMPull to distinguish between disease-derived and control aggregate-containing samples according to tau aggregate abundance and brightness.

### Super-resolved SiMPull quantifies aggregate morphology with single particle precision

To characterize aggregate morphology with a precision unattainable via diffraction-limited imaging, we adapted the SiMPull assays for Stochastic Optical Reconstruction Microscopy (STORM)^24^. We were then able to quantify the length, perimeter, total area, and eccentricity of individual aggregates in brain homogenate. We observed two dominant aggregate shapes, loosely falling into extended ‘fibrillar’ and short ‘globular’ categories (Fig. 3A and B and Supplementary Fig. 2A). As anticipated, aggregates were highly heterogenous, ranging from 30 nm to more than 400 nm in length. On average, tau aggregates detected in AD brain were 115 nm long while the ones detected in control brains were only 95 nm long (*p* = 0.064). While we did not observe any significant differences in the mean size or shape of the aggregates (Fig. 3C), we did observe qualitative separation of the cumulative distribution of aggregate length and eccentricity between disease-derived and control samples (Fig. 3D). This prompted us to consider the proportion of long aggregates (>250 nm), which was on average 1.5× higher in the AD samples compared to control (Fig. 3E, t-test *p* = 0.045). Similarly, the proportion of fibrillar aggregates (eccentricity >0.9) was 25% in AD while only 20% in the control brains (Fig. 3E, t-test *p* = 0.0072). This suggests that the accumulation of aggregates with specific sizes and shapes may be correlated with disease pathology.

**Fig. 3.**
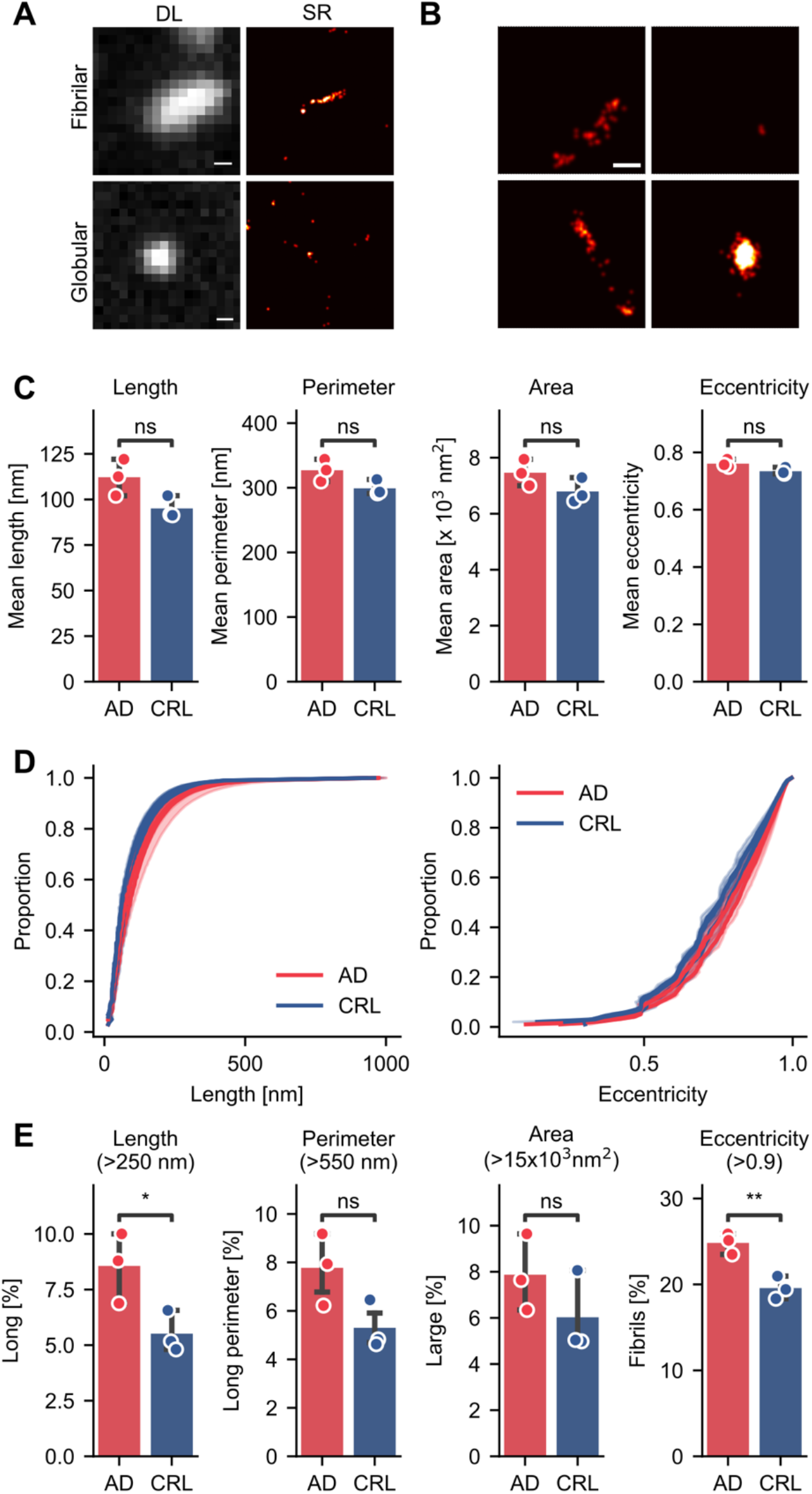
Characterizations of brain-derived tau aggregates using SiMPull coupled to super-resolution microscopy. **(A)** Representative images of diffraction-limited and super-resolved tau aggregates revealing distinct morphological categories. Scale bar = 200 nm. **(B)** Example images of aggregates of varying size and eccentricity. Scale bar = 100 nm. **(C)** Mean length, perimeter, area, and eccentricity of p-tau aggregates detected via AT8 SiMPull. **(D)** Cumulative distribution of aggregate length and eccentricity measured as in C. **(E)** Proportion of aggregates satisfying various thresholds for size (length, perimeter, area) and shape (eccentricity) for brain-derived samples taken from AD (red) and control (blue) cohorts. Panel C and E show the mean ± S.D. of n = 3 biological replicates using a t-test. ns: *p* > 0.05, *: *p* < 0.05, **: *p* < 0.01.

The intersection of size and shape for individual aggregates provides another metric for comparing aggregate populations between cohorts (Supplementary Fig. 2B and C). In AD and control samples, longer aggregates (> 250 nm) were significantly more likely to be fibrillar; 55% of long aggregates had an eccentricity > 0.9, compared to 5% of short (< 100 nm) aggregates (Supplementary Fig. 2D, AD *p* = 0.000018). Interestingly, the percentage of long aggregates in the population of round aggregates (eccentricity <0.7) was significantly higher in AD than in control (AD: 2.5%, CRL: 1%, *p* = 0.047) (Supplementary Fig. 2E). Overall, this shows that tau aggregates with a wide variation in size and shape are formed in both control and AD brain. Importantly, changes in the morphology of aggregate populations may be an important indicator of disease pathology and progression in AD.

Our resolution did not allow aggregates <50 nm to be identified as fibrils (with eccentricity >0.9). We confirmed that when filtering out all aggregates <50 nm in length, we still see a significant difference in the proportion of long aggregates and fibrils between the brains from AD and control patients, as before (Supplementary Fig. 2F). Additionally, we observed a significant difference in the mean length, eccentricity, and perimeter of aggregates between AD and control samples once aggregates < 50 nm had been removed (Supplementary Fig. 2G). This may indicate that the variability between patients within a cohort is driven by small aggregates <50 nm, such that once these are removed the mean values are more consistent within disease cohorts (coefficients of variation: 9% and 5% in AD, 6% and 2% in CRL, before and after filtering < 50 nm respectively).

### Aggregate composition is accessible via co-labelling of multiple epitopes

We next sought to characterize the composition of tau aggregates by adapting the SiMPull assay for co-localization studies. This enabled us to detect several phosphorylation sites present in the same aggregates simultaneously, namely p-Ser202 and p-Thr205 (AT8), and p-Thr181 (T181). We captured tau aggregates either with AT8 or T181 antibodies and then detected with a mixture of both the AT8 and T181 antibodies labelled with spectrally distinct fluorophores (AlexaFlour 647 and 488 respectively, Fig. 4A). These configurations ensure that we capture aggregates (i.e., particles containing two copies of the capture antibody epitope) decorated with the modifications of interest.

**Fig. 4.**
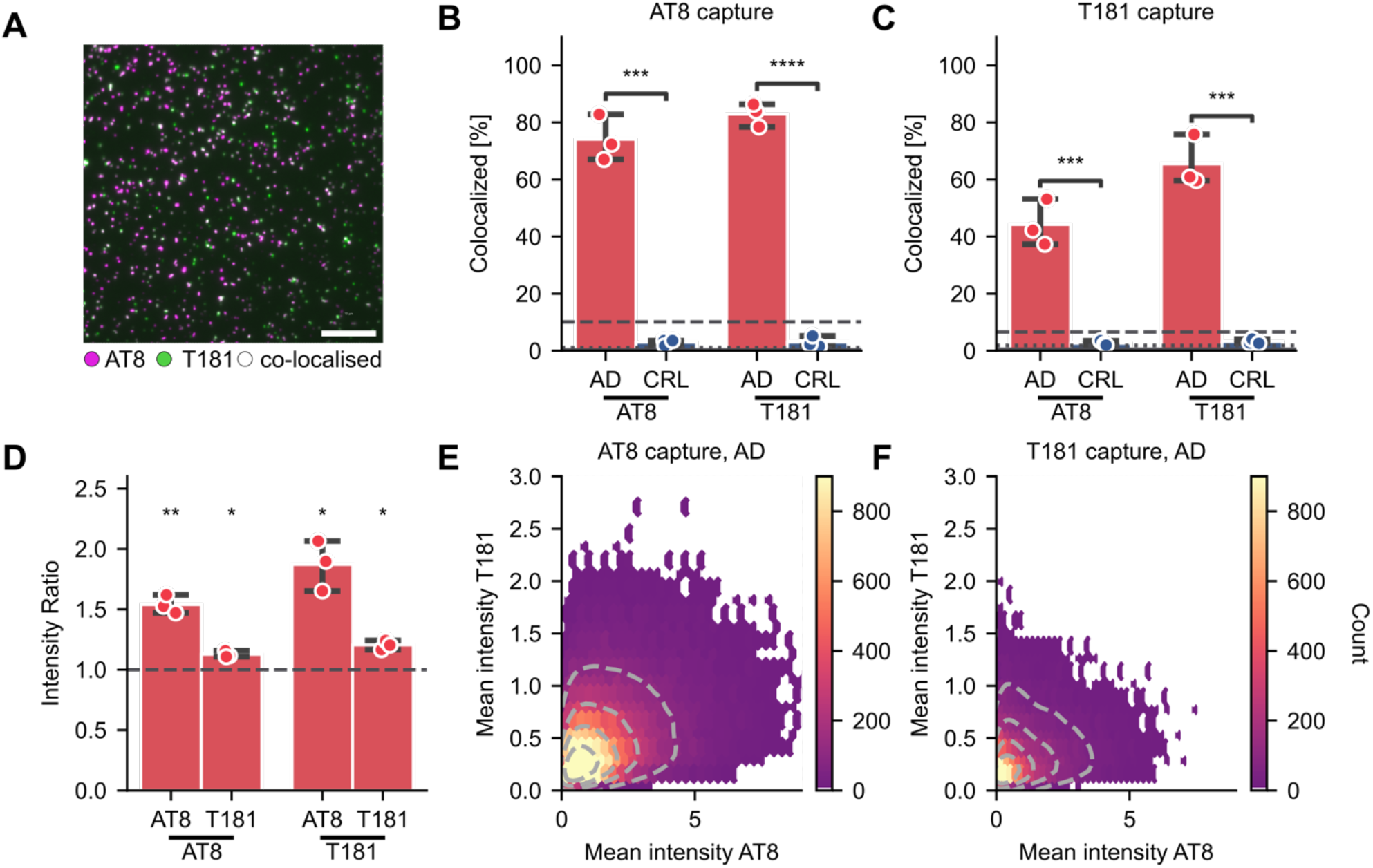
Co-localization of antibodies targeting tau phosphorylation sites (T181, AT8) reveals differences in disease-associated aggregate composition. **(A)** Representative image of co-localized spots in AD brain homogenate. Scale bar = 10 μm. **(B-C)** Percentage of green spots (T181) co-localized with magenta spots (AT8) and vice versa using **(B)** AT8 or **(C)** T181 to capture aggregates from brain homogenate. **(D)** Ratio of the brightness of co-localized and non-colocalized spots for each channel in B, C. (AT8: 638 nm, T181: 488 nm) detected in AD samples using either AT8 or T181 for capture. **(E-F)** Mean intensity of each co-localized spot in AD samples using **(E)** AT8 or **(F)** T181 to capture. Panel B and C show the mean ± S.D. of n = 3 biological replicates in each disease cohort, compared by t-test. *: *p* < 0.05, **: *p* < 0.01, ***: *p* < 0.001. Panel D: one-sample t-test against hypothetical value of 1 (equivalent to no difference between the co-localized and non-colocalized spots).

To quantify co-localization, we detected diffraction-limited spots labelled by one fluorophore then calculated the proportion of those spots that were also labelled with the second fluorophore. Almost no co-localization of the T181 and AT8 antibodies was observed in control samples (no more than would occur by chance; Fig. 4B dotted line, obtained from rotating one channel relative to the other). Comparatively, in the AD sample more than 75% of T181-positive tau aggregates were also positive for AT8, and vice versa (Fig. 4B), significantly higher than in the control samples (t-test: AT8 detection *p* = 0.00011, T181 detection *p* = 0.0000066). Comparable observations were made using T181 to capture tau aggregates (Fig. 4C, t-test: AT8 detection *p* = 0.00090, T181 detection *p* = 0.00028). Finally, we combined the co-localization and brightness analyses. We found co-labelled aggregates were significantly brighter than singly labelled aggregates in either SiMPull configuration in both detection colors (Fig. 4D, one-sample t-test, AT8-AT8: *p* = 0.0065, AT8-T181: *p* = 0.014, T181-T181: *p* = 0.011, T181-AT8: *p* = 0.018). Indeed, co-labelled aggregates were up to twice as bright (∼large) in comparison to singly labelled aggregates in the AD brain samples. Lastly, we observed a significant positive correlation (AT8 capture Pearson’s coefficient = 0.36, T181 capture: Pearson’s coefficient = 0.28) in the brightness of individual aggregates between channels indicating that, despite being heterogeneous across the population, aggregates with more AT8 phosphorylation sites also tend to have increased T181 phosphorylation (Fig. 4E and F). Overall, these data demonstrate that SiMPull is amenable to compositional studies of tau aggregates that reveal additional discriminators between AD and control cohorts.

### SiMPull reveals heterogeneity among tau aggregates derived from tissue versus biofluids

We lastly confirmed our assay is compatible with readily available and clinically relevant samples, such as serum. For this purpose, we tested 9 human serum samples obtained from AD patients and 9 from healthy control patients. We detected significantly more HT7-positive spots in human serum samples compared to a tau-negative BSA control (Fig. 5A, one-way Anova: *p* = 0.00020; post-hoc Tukey HSD test, *p* = 0.00064 for CRL vs BSA, *p* = 0.00013 for AD vs BSA). This confirmed the compatibility of the tau SiMPull assays for use with human serum. We were able to detect an average of 770 ± 160 HT7-positive spots per FOV in the AD serum and 670± 170 in the control serum. However, there was no significant difference in the number of spots in serum from AD versus control patients (*p* = 0.44). We did not observe any correlation to the age and sex of the donors.

**Fig. 5.**
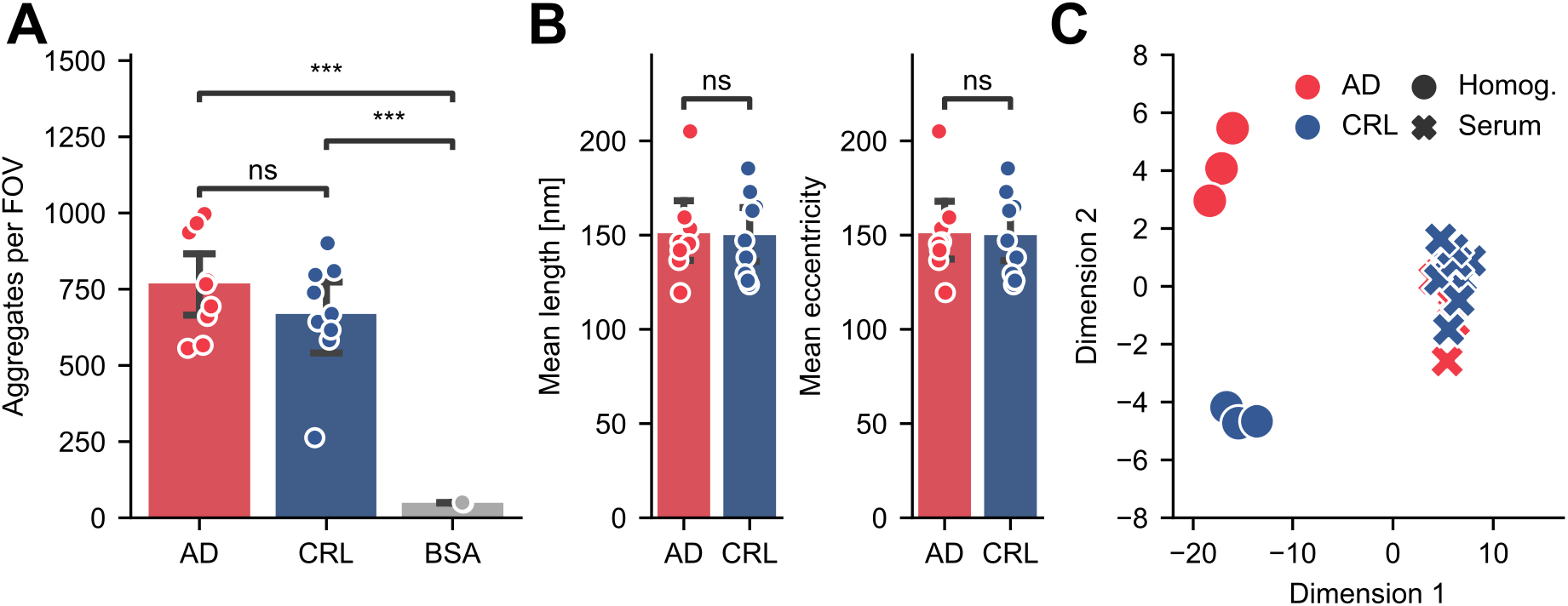
Quantification of total tau aggregates in human serum. **(A)** SiMPull quantification of HT7-positive aggregates in human serum samples from AD and control donors, and a BSA negative control. **(B)** Mean length and area in serum tau aggregates determined by super-resolution microscopy. **(C)** Linear discriminant analysis (LDA) of human serum (n = 9) and brain homogenate (n = 3) samples from AD and control patients. Panel A and B show the mean ± S.D. of n = 9 biological replicates (BSA n = 2 technical replicates). Panel A reports a one-way Anova with post-hoc Tukey HSD test, Panel B reports Student’s t-tests. ns: *p* > 0.05, **: *p* < 0.01, ***: *p* < 0.001.

In order to characterize any morphological differences associated with disease we also super-resolved the serum-derived aggregates. We were unable to detect significant differences in either length or eccentricity between the HT7-positive tau aggregate population in AD compared to control samples (Fig. 5B). Proportional analyses as described above for AT8-positive brain-derived aggregates are certainly possible. However, we reasoned that the combination of morphological information may better encapsulate subtle differences in the aggregate populations both between tissue types and disease cohorts. We compiled the mean morphological parameters collected for each aggregate population, including length, area, perimeter, and eccentricity supplemented with the number of aggregates, number of localizations per aggregate, and major and minor axis length. A pairwise comparison of all the features is shown in Supplementary Fig. 3. These parameters were used for linear discriminant analysis (LDA), a method commonly used for supervised dimensionality reduction which maximizes the separation between different categories using linear combinations of the provided parameters.

Using this method, it was possible to examine whether the HT7-positive aggregates detected in serum differed from those observed in human brain homogenate, as well as any differences between AD and control cohorts. Thus, the dataset included four different aggregate populations; namely, HT7-positive aggregates from control or AD serum, and from control or AD brain homogenate. Interestingly, when projected onto a two-dimensional parameter space we observed the sample cohorts formed three clusters driven primarily by eccentricity, minor and major axis length (Fig. 5C). Brain homogenate and serum samples were well separated by this method, clustering to opposite sides of the first dimension. This indicates that the tau aggregates in serum are distinguishable from the brain-derived aggregates according to the parameters collected from the HT7 SiMPull assay. In addition, AD and control brain-derived samples formed two distinct clusters separated in the second dimension, indicating aggregates from these samples also have distinguishing morphological features captured by the HT7 SiMPull analysis. However, no separation between the aggregates from AD and control serum was observed in this dimension. This is consistent with recent studies which showed that tau found in blood mostly originates from peripheral tissues and is not brain derived^25^, resulting in a lack of correlation between serum and CSF total tau^26^. If the majority of tau aggregates in serum are also composed of non-brain-derived tau, it is perhaps unsurprising that we do not detect differences in this population associated with neurodegenerative disease.

## Discussion

We describe here a SiMPull assay that enables the detection and characterization of tau aggregates in a variety of biological samples with high specificity and sensitivity. This assay produces complementary readouts of aggregate number, morphology, and composition. For this study, we focused on two commonly used antibodies, HT7 and AT8, to detect total tau aggregates and p-tau aggregates respectively, as well as an additional phospho-tau marker T181. We anticipate this method will be readily applicable to other tau aggregate species from various biological samples using antibodies targeting a range of characteristics including post-translational modifications, conformation, and isoforms ^27–29^.

Notably, we found that the simple number of aggregates is not necessarily sufficient to discern between disease and control samples. Given the stark differences between insoluble aggregate accumulation in health and disease on which Braak staging is predicated, this was unexpected. However, SiMPull is able to detect small, soluble aggregates which are invisible to many techniques. Their comparable abundance in disease and age-matched healthy brains suggests differences in aggregates driving disease must lie beyond their abundance. The groups’ differentiation was aided by additional features which can be read out from standard SiMPull (brightness ≈ size) and SiMPull extended via super-resolution or co-localization analyses (size and composition). Using these characteristics, it was possible to detect significant differences between tau aggregates derived from AD and control human brain homogenate samples. Specifically, we observed an increased proportion of large aggregates in brain homogenate from AD patients. Tau aggregates in AD samples were also more likely to be decorated by multiple post-translational modifications as revealed by our co-localization studies. Those studies could be expanded in the future by determining the size of the co-localized aggregates to assess whether the aggregates are indeed bigger (i.e., contain more total tau) or the proportion of phosphorylated sites is increased. High variability between AD-derived samples limited the ability to make statistical inferences in some cases, and future studies designed to specifically compare AD with control cohorts should consider larger cohort sizes.

Using the tau SiMPull assay, it is possible to detect single aggregates and characterize them individually, enabling the identification of subpopulations which are potentially pathologically relevant. This is not possible with bulk techniques such as ELISA which derive the average of a characteristic across the entire population. Importantly, an overall increase in tau phosphorylation as well as changes in the tau phosphorylation pattern have been linked to progression in AD. Levels of phosphorylated tau in cerebrospinal fluid (CSF) and blood are a core biomarker for AD^30^, and elevated levels of pThr181 have been detected in the pre-clinical stage of AD suggesting its potential for early diagnosis^31^. Yet, the assays detected all tau, including tau monomers, and little was known about tau aggregates in clinical samples. In the brain, high heterogeneity in the frequency and occurrence of tau phosphorylation between patients has been observed^32,33^. While certain sites (mostly towards the N and C termini) are phosphorylated only in the disease state, phosphorylation of other sites also occurs in healthy adult human brains but with much lower prevalence compared to AD^32^. Other sites, e.g. T231, show a progressive increase in phosphorylation with disease state^34^. However, studies so far have only analyzed aggregate populations using methods unable to distinguish individual particles (ELISA, mass spectrometry), giving an average of the phosphorylation sites of the entire aggregate population. Thus, little is known about the phosphorylation status of individual aggregates and a potentially causal relationship between certain modifications.

Measuring the composition of the individual aggregates, i.e., the occurrence of several phosphorylation sites in a single aggregate, is not readily possible with bulk techniques. While other single-molecule methods to study tau aggregates exist which report a digital assay readout of diffraction-limited pixel positivity, it is not amenable to characterizing aggregate size or shape^35^. The results from our study indicate that those features might play a greater role in disease pathology compared to just aggregate abundance, as we found significantly more large tau aggregates in AD brains. Interestingly, we observed a greater difference between aggregate brightness obtained through the diffraction-limited assay than in aggregate length between AD and control. Their two-dimensional nature means the super-resolved morphological parameters are potentially impacted by aggregate orientation relative to the *xy* plane, while the brightness we expect to be agnostic. However, due to the characteristics of STORM where only a small (unknown) fraction of fluorophores is being turned on, it is difficult to directly compare diffraction-limited measurements with super-resolved ones^36^. Regardless, SiMPull is sufficiently sensitive to detect small soluble tau aggregates which cannot be readily detected by many of the established techniques. Thus, it may be possible to use tau SiMPull for the early detection and monitoring of tau pathology as it spreads through the brain ahead of the formation of large insoluble aggregates.

Importantly, we have not only observed changes in fibrillar aggregates but also round ones. Brain-derived round aggregates appear to be longer in AD compared to control cohorts. This is in line with the analysis of the free-energy landscape showing two pathways of tau aggregation, one leading to ordered fibrils and the other to amorphous phases^37^. Furthermore, spontaneous aggregation of *in vitro* hyperphosphorylated tau forms round aggregates which were able to induce an inflammatory response in human macrophages^38^. This suggests that both fibrillar and hyperphosphorylated round tau aggregates may be associated with disease pathology, and tau SiMPull assays readily enable their distinction in biological contexts.

Overall, the specificity and sensitivity of the tau SiMPull coupled with its ease of use and low sample volume requirements (< 10 μL) make it perfectly suited to characterize model systems or clinical samples. This technology makes temporal monitoring of such systems feasible, producing longitudinal data with which it would be possible to tackle fundamental mechanistic questions such as the order of phosphorylation before and after aggregate formation, or whether phosphorylation at particular sites triggers hyperphosphorylation. Further, the compatibility of tau SiMPull with readily accessible biofluids coupled with the ability to distinguish between disease-derived samples from matched control cohorts promises to revolutionize diagnostic and disease monitoring strategies.

## Methods

### Materials

Post mortem brain tissue from three donors with AD and three age-matched control donors was acquired from the Cambridge Brain Bank (with the approval of the London—Bloomsbury Research Ethics Committee; 16/LO/0508, Supplementary Table 1; and with written informed consent from either the donor ante mortem or from their next of kin post mortem in accordance with UK law). The brain samples were voluntarily donated without any compensation. Serum samples from ten people with amnestic Alzheimer’s disease (with dementia or mild cognitive impairment supported by imaging and/or biomarker evidence of AD) and ten controls were provided ante mortem after written informed consent from volunteers with mental capacity (with the approval of East of England Cambridge Central Research Ethics Committee 15/EE/0270; see Supplementary Table 2).

### Preparation of recombinant aggregates

Lyophilized monomeric recombinant Aβ42 peptide (Stratech, Cat. No. A-1170-2-RPE-1.0mg) was dissolved in PBS (pH = 7.4) at 200 μM on ice. The solution was quickly aliquoted and snap frozen. To prepare recombinant Aβ42 fibrils, an aliquot was thawed and diluted to 4 μM in 1xPBS supplemented with 0.01% NaN3 (Merck, Cat. No. 71290) and incubated at 37 °C under quiescent conditions for one week. The Aβ42 fibril was then sonicated as described previously^39^ with modification. The one-week aggregated Aβ42 aliquot was immersion sonicated in an ice water bath with a 3-mm-titanium probe (Sonicator microprobe 4422, Qsonica) mounted on a tip sonicator (Ultrasonic processor Q125, QSonica) at 20 kHz with 40% of power for 24×5-s bursts with 15-s rests between bursts. Thereafter, the sonicated aggregate was centrifuged, aliquoted (50 μL) and snap frozen. The aliquots were stored at -80 °C until use.

Wild type α-synuclein was expressed, purified in *E. coli* and stored at -80 °C as described previously^40^. To remove pre-aggregation seeds, the solution was ultracentrifuged at 91,000 g at 4 °C for 1 hour (Optima TLX Ultracentrifuge, Beckman). The concentration of the supernatant was then determined by A280 (ε280 = 5,960 M−1 cm−1). The supernatant was then diluted to 70 μM in 1xPBS supplemented with 0.01% NaN3 and incubated at 37 °C with shaking at 200 rpm for two months.

Recombinant N-terminally 6xHis-tagged human P301S 0N4R tau was expressed and purified from *E. coli* BL-21 DE3 cells. Protein expression was induced by addition of 0.5 mM IPTG at 16 °C overnight. Cells were pelleted (17,000 x g, 3 min) and lysed in recombinant tau-lysis buffer (1 mM benzamidine, 1 mM PMSF, 1x cOmplete™, EDTA-free Protease Inhibitor Cocktail mix (Merck), 14 mM b-mercaptoethanol, 300 mM NaCl, 25 mM HEPES, 30 mM imidazole, 1% NP-40). Purification was performed on the AKTA Pure using the HisTrap HP column (Cytiva), followed by size exclusion chromatography using a Superdex 200 HiLoad 16/600 pg column as previously described (PMID: 29768203). Tau monomer fractions were stored in 1x PBS buffer, freshly supplemented with 1 mM DTT. *In vitro* aggregated tau assemblies were prepared by addition of heparin at 37 °C for 3 days while shaking, using monomeric tau at 60 μM in the presence of 20 μM heparin (Sigma Aldrich) in PBS supplemented with 2mM DTT and 1x cOmplete™, EDTA-free Protease Inhibitor Cocktail mix. A small aliquot of the assemblies was kept for analysis and the remaining material was sonicated for 15 sec before long-term storage at -80 °C.

The HT7 epitope-containing peptides were designed with the following sequences GAA**PPGQK**GQ{PEG4}GAA**PPGQK**GQ to mimic dimeric particles and GAA**PPGQK**GQ{PEG4}GKPQPAGAQG to mimic monomeric particles, where PPGQK corresponds to the immunogen’s amino acid sequence (HT7: abcam, Cat. No. MN1000). These peptides were synthesized by GenScript and provided as lyophilized powder before being resuspended in milliQ water at a final concentration of 2 mg/mL. Stocks were stored at -20 °C, then diluted to the desired concentration in PBS immediately before use.

### Isolation of cell-derived tau aggregates

HEK293 cells expressing tau P301S-Venus (PMID: 28049840) were maintained in DMEM supplemented with 10% FCS, 100 U/ml penicillin, 100 ug/ml streptomycin and grown at 37 °C and 5% CO2. The cells were seeded with 50 nM heparin-assembled recombinant 6xHis-tau P301S assemblies in the presence of 1% Lipofectamine2000 and by serial dilutions, a single clone was isolated (R1E5) that stably propagates tau P301S-Venus aggregates. Cells were lysed in R1E5-Lysis buffer (1x PBS, 1% w/v Triton X-100, 1x cOmplete™, EDTA-free Protease Inhibitor Cocktail mix, 1x PhosSTOP™ phosphatase inhibitor mix) on ice for 30 min. The lysate was subsequently centrifuged at 14, 000 × g for 15 min at 4 °C and the clarified lysate was aliquoted and stored at -20 °C.

### Homogenization of Brain Samples

Brains were flash-frozen and stored at −80 °C at the Cambridge Brain Bank in Cambridge. Fresh-frozen brain tissue was homogenized using a method adapted from Goedert, et al.^41^. Briefly, the tissue was homogenized at 4 °C in a VelociRuptor V2 Microtube Homogenizer (Scientific Laboratory Supplies, Cat. No. SLS1401) in 10 volumes of homogenization buffer (10 mM Tris-HCl, 0.8 M NaCl, 1 mM EGTA, 0.1% Sarkosyl, 10% sucrose; pH 7.32) containing cOmplete™ Ultra Protease Inhibitor and PhosStop™ Phosphatase Inhibitor. The homogenate was centrifuged at 21,000 × g for 20 minutes at 4 °C, and the upper 90% of the supernatant was retained. The pellet was re-homogenized in 5 volumes of homogenization buffer, then centrifuged at 21,000 × g for 20 minutes at 4 °C. The upper 90% of this supernatant was then removed and combined with the first supernatant, and this mixture was aliquoted and frozen at -80 °C until used for further experiments. Total protein concentration was determined using a BCA assay (Thermo Fisher, Cat. No. 23227) as per the manufacturer’s instructions. The concentration of tau was determined using the Human Tau ELISA kit (abcam, Cat. No. ab273617) according to the manufacturer’s instructions.

### Coverslip passivation

Coverslip passivation for SiMPull was completed using a method adapted from Chandradoss, et al.^42^. 26 mM × 76 mM #1.5 borosilicate glass coverslips (VWR, Cat. No. MENZBC026076AC40) were washed in a sonication bath for 10 minutes each in 18.2-MΩ·cm water, acetone, and methanol, respectively, then 20 minutes in 1 M KOH. Coverslips were rinsed with 18.2-MΩ·cm water and then methanol, then dried with nitrogen, then cleaned with argon plasma for 15 minutes (Femto Plasma Cleaner; Diener Electronic, Royal Oak, MI, USA). Next, coverslips were silanized in a 3:5:100 mixture of (3-Aminopropyl)triethoxysilane (APTES) (Fisher Scientific UK, Cat. No. 10677502), acetic acid, and methanol, respectively. Coverslips were sonicated in this solution for 1 minute, left undisturbed for 10 minutes, sonicated again for 1 minute, then left undisturbed for an additional 10 minutes. Coverslips were then rinsed twice with 18.2-MΩ·cm water and once with methanol, then dried with nitrogen. One 50-well silicone gasket (Grace Bio-Labs, SKU 103250) was then attached to the surface of each coverslip. Each well was passivated by firstly introducing 9 μL of a freshly prepared 100:1 aqueous mixture of SVA-PEG-OMe (110 mg/mL, Mw ∼5,000; Laysan Bio Inc., Cat. No. MPEG-SVA-5000) and SVA-PEG-Biotin (100 mg/mL, Mw ∼5,000; Laysan Bio Inc., Cat. No. Biotin-PEG-SVA-5000) followed by adding 1 μL of 1 M NaHCO3 (pH 8.5). Coverslips were left covered in a humidity chamber for 24 hours, then rinsed twice with 18.2-MΩ·cm water, then dried with a nitrogen stream. Each well was then further passivated by adding 9 μL of a freshly prepared aqueous solution of methyl-PEG4-NHS-Ester (10 mg/mL, Thermo Fisher, Cat. No. 22341), followed by adding 1 μL of 1 M NaHCO3 (pH 8.5). Coverslips were left covered in a humidity chamber for 24 hours, then rinsed twice 18.2-MΩ·cm water, dried with nitrogen, and then stored desiccated at -20 °C until used for experiments.

### Antibody Labelling

Biotin-conjugated AT8 and HT7 are commercially available (AT8: Invitrogen, Cat. No. MN1020B; HT7: Invitrogen, Cat. No. MN1000B). T181 (abcam, Cat. No. ab232849 and SC211 (Santa Cruz, Cat. No. sc-12767L) were biotinylated site-specifically through Click chemistry, using the SiteClick Antibody Azido Modification kit (Invitrogen, Cat. No. S20026). Briefly, 200 μg of the antibody was buffer-exchanged and concentrated to ∼2 mg/ml and subsequently incubated with β-galactosidase overnight. The antibody was then incubated overnight in the presence of β-1,4-galactosyltransferase to attach azide modified carbohydrates (UDP-GalNAz) to the modified glycan chains. Subsequently, the antibody was purified using a 50 kDa Amicon Ultra spin column (Merck). Ten equivalents of DBCO-PEG4-biotin (Sigma Aldrich, Cat. No. 760749) were added to the antibody and incubated overnight at 37 °C. The biotin-labelled antibody was purified using a 50 kDa Amicon Ultra spin column, and the concentration was quantified by A_280_. Antibodies were fluorescently labelled with AF647 or AF488 using the Alexa Fluor® Conjugation Kit (Fast) - Lightning-Link (Abcam, Cat. No. ab269823 and ab236553) according to manufacturer’s instructions. Excess fluorophore was removed using a 50 kDa Amicon Ultra Spin column (Merck) and a 40 kDa Zeba Spin desalting column.

### Single-molecule Pulldown (SiMPull)

SiMPull for tau aggregates was adapted from^43^. Briefly, wells were washed once with PBST (PBS (50 mM tris base and 150 mM NaCl, pH 7.4) containing 0.5% Tween-20). 10 μL NeutrAvidin (0.2 mg/mL in PBST; Thermo Fisher, Cat. No. 31000) was added to each well and left to incubate for 10 minutes. After a washing sequence (two 10-μL washes of PBST, followed by one 10-μL wash of PBS containing 1% Tween-20), biotinylated capture antibodies (AT8: Invitrogen, REF MN1020B; HT7: Invitrogen, REF MN1000B; T181: ab232849 were diluted to 10 nM in blocking solution (1 mg/mL BSA in PBST), and left to incubate for 10 minutes. After a washing sequence, wells were blocked with blocking solution for 10 minutes. After one PBST wash, 10 μL of sample was added to each well and left to incubate. Brain homogenate samples were diluted 1:10 in PBS and incubated for 1h at room temperature, serum samples were diluted 1:2 in PBS and incubated overnight at 4 °C, followed by a washing sequence. Fluorescently labelled detection antibodies (AT8: Invitrogen, Cat. No. MN1020, 5 nM; HT7: Invitrogen, Cat. No. MN1000, 2 nM; T181: abcam, Cat. No. ab232849, 5 nM) were added in blocking solution, and left for 15 minutes, followed by a final washing sequence. Wells were then washed once with PBS, then the wells were filled with fresh PBS before the gasket was sealed with another clean coverslip.

### Cross-reactivity of tau SiMPull assay

AT8 and HT7 SiMPull assays were performed as described above, using recombinant α-synuclein and amyloid-β aggregates at 1 μM. As a control, SiMPull was performed using amyloid-β (6E10, BioLegend, Cat. No. 803007 and 803021) and α-synuclein (SC211, Santa Cruz, Cat. No. sc-12767) specific antibodies for matched capture and detection.

### Diffraction-limited Imaging and Analysis

Imaging was done on a home-built total internal reflection fluorescence (TIRF) microscope, consisting of an inverted Ti-2 Eclipse microscope body (Nikon) fitted with a 1.49 N.A., 60x TIRF objective (Apo TIRF, Nikon) and a perfect focus system. Images were acquired using a 638 nm laser (Cobolt 06-MLD-638, HÜBNER) or 488 nm laser (Cobolt 06-MLD-488, HÜBNER). Detection antibodies labelled with AF647 or AF488 were excited at 638 nm and 488 nm respectively, with the resultant fluorescence collected by the objective, passed through a quad-band dichroic beam splitter and cleaned up using the appropriate emission filter (for 488-nm-induced fluorescence: BLP01-488R-25x36, Semrock and FF01-520/44-25x36, Semrock; for 638-nm-induced fluorescence BLP01-635R-25x36, Semrock).

EMCCD camera (Evolve 512, Photometrics) operating in frame-transfer mode (electron-multiplying gain of 6.3 electrons/ADU and 250 ADU/photon). Each pixel corresponds to a length of 107 nm on the recorded image.

Typically, 16 fields of view (FOVs) of 54.784 μm^2^ each were imaged per well, each for 50 frames of 50 ms exposure. Each pixel corresponds to a length of 107 nm. Images were collected in a grid using an automated script (Micro-Manager^44^) to avoid any bias in the selection of FOVs. For co-localization experiments, images were acquired sequentially in each excitation channel.

Individual fluorescent spots were quantified using a python-based adaptation of ComDet^45^, a package originally developed to identify bright intensity spots in images with a heterogeneous background. Briefly, a mean intensity projection was prepared from the last 40 frames of each field of view to minimize background fluctuations. Thresholds were then optimized for each experiment by comparing the number of particles identified by ComDet in positive and negative control images, such that the threshold was set to the lowest value at which the negative control was less than 1% of the positive control. This threshold was then used along with a particle size estimate of 4 to identify fluorescent spots. The mean intensity of each spot was measured using scikit-image, and background corrected by subtracting the median intensity of all non-spot pixels for each FOV.

For co-localization experiments, ComDet identification was performed independently on both detection channels and the resultant list of centroid coordinates collected. For each channel, spots co-labelled in the opposing channel were determined as those within a 4-pixel radius according to the Euclidean distance (Supplementary Fig. 4A). In the case where more than one spot met this criterion, the closest spot was selected as the co-localized pair. The chance co-localization was estimated by transposing the centroid coordinates for the second channel in the x-dimension and repeating the threshold distance analysis (Supplementary Fig. 4B). Finally, the proportion of co-localized spots was calculated for a given channel as the number of spots matched to a corresponding spot divided by the total number of spots detected in that channel (Supplementary Fig. 4C).

### Super-resolution Imaging and Analysis

STORM buffer was freshly prepared at 1 mg/mL glucose oxidase, 52 μg/mL catalase and 50 mM MEA (cysteamine) in 50 mM Tris in PBS + 10% Glucose, pH 8, and filtered through a 0.02 μm filter. In preparation for STORM imaging, PBS was removed from each well and replaced with STORM buffer before sealing the gasket with another clean coverslip. Imaging was performed with Typically, at least 6000 frames of 33 ms exposure were recorded per field of view.

Super-resolution images were reconstructed using the Picasso^46^ package. Briefly, after discarding the first 300 frames, localizations were identified and fit then corrected for microscope drift using the inbuilt implementation of redundant cross-correlation. Localizations were then filtered for precision <30 nm, with a final mean precision of ≈ 12 nm. Random localizations were removed using DBSCAN as provided by the scitkit-learn package^47^ with permissive parameters (radius of 2 and minimum density of 1). The resultant clustered array of localizations was then subjected to a series of morphological dilation, closing and erosion operations as provided by the scikit-image package^48^ to yield single connected regions of interest corresponding to individual aggregates (Supplementary Fig. 5 A - C). Aggregates with <3 localizations per cluster were considered noise and removed. Each aggregate was then measured using a combination of scikit-image^48^ for basic region properties such as perimeter, area, and eccentricity, and SKAN^49^ for skeletonized length (Supplementary Fig. 5D) whereby the length of each aggregate is reported as the summed branch distance. Finally, super-resolved images were rendered using the inbuilt Picasso^46^ functionality.

### Limit of Detection

The limit of detection (LOD) was determined by a serial dilution of a brain homogenate sample obtained from an AD donor. The absolute concentration of tau was determined using the commercially available human tau ELISA. It should be noted that the ELISA was not aggregate specific, thus it reflects the total concentration of all tau (including monomer) in the sample. SiMPull were performed and the resultant particle counts at each homogenate concentration were fit using four-parameter logistic regression (Supplementary Fig. 1C).

The limit of blank (LOB) is the highest apparent number of spots expected to be found when replicates of a sample containing no tau aggregates are detected, and is defined as^23^:

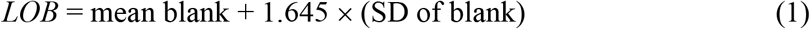

The limit of detection (LOD) was determined by utilizing the previously determined LOB and replicates of a sample containing a low concentration of tau aggregates, and is given by the expression ^23^:

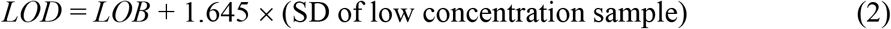

The LOB for the HT7 assay was determined as 575 pg/mL and the LOD as 874 pg/mL. The LOB for the AT8 assay was determined as 591 pg/mL and the LOD as 2201 pg/mL.

### Statistical analysis

Statistical analyses were performed using either the SciPy^50^ or statsannotations^51^ packages in python. The exact *p* values and statistical details are provided in the main text or figure legends as appropriate.

### Data availability

Where feasible, exemplar images for diffraction-limited and super-resolved analyses have been provided alongside summaries of the pre-processed quantitative data via Zenodo DOI: 10.5281/zenodo.8020036. Any other data supporting this study are available from the corresponding author upon reasonable request.

### Code availability

Analyses presented in this manuscript rely on various published python packages including Picasso^46^, scikit-learn^47^, scikit-image^48^, and SKAN^49^. All other custom python scripts used in this study are available via Zenodo DOI: 10.5281/zenodo.8027256.

## Acknowledgments

We thank Yunzhao Wu for technical assistance and Emre Fertan for helpful discussion. This work was supported by the UK Dementia Research Institute, which receives its funding from DRI Ltd., funded by the UK Medical Research Council, Alzheimer’s Society, and Alzheimer’s Research. D.C. was supported by the Lady Edith Wolfson Junior Non-clinical Research Fellowship awarded by the MND Association UK (Cox 971-799). J.Y.L.L. is supported by the Croucher Foundation Limited (Hong Kong). W.A.M and T.K. received funding from the Innovative Medicines Initiative 2 Joint Undertaking under grant agreement 116060 (IMPRiND). This Joint Undertaking receives support from the European Union’s Horizon 2020 Research and Innovation Program and EFPIA. This work is supported by the Swiss State Secretariat for Education, Research, and Innovation (SERI) under contract 17.00038. JBR is supported by the Wellcome Trust (103838; 220258), the Medical Research Council (MC_UU_00030/14); JBR and the Cambridge Brain Bank are supported by the NIHR Cambridge Biomedical Research Centre (NIHR203312). The views expressed are those of the authors and not necessarily those of the NIHR or the Department of Health and Social Care. For the purpose of open access, the authors have applied a CC BY public copyright licence to any Author Accepted Manuscript version arising from this submission.

## Author contributions

Conceptualization: DK, DB

Formal Analysis: DB, DC

Funding acquisition: DK

Methodology: DB, DC, MB

Investigation: DB, MB

Resources: JYLL, TK, WAM, JBR

Software: DC, DB

Supervision: DK

Visualization: DB, DC

Writing—original draft: DB, DC

Writing—review & editing: DK, DB, DC, MB, JYLL, TK, JSHD, JBR, WAM

## Supplementary Materials for

**Supplementary Fig. 1.**
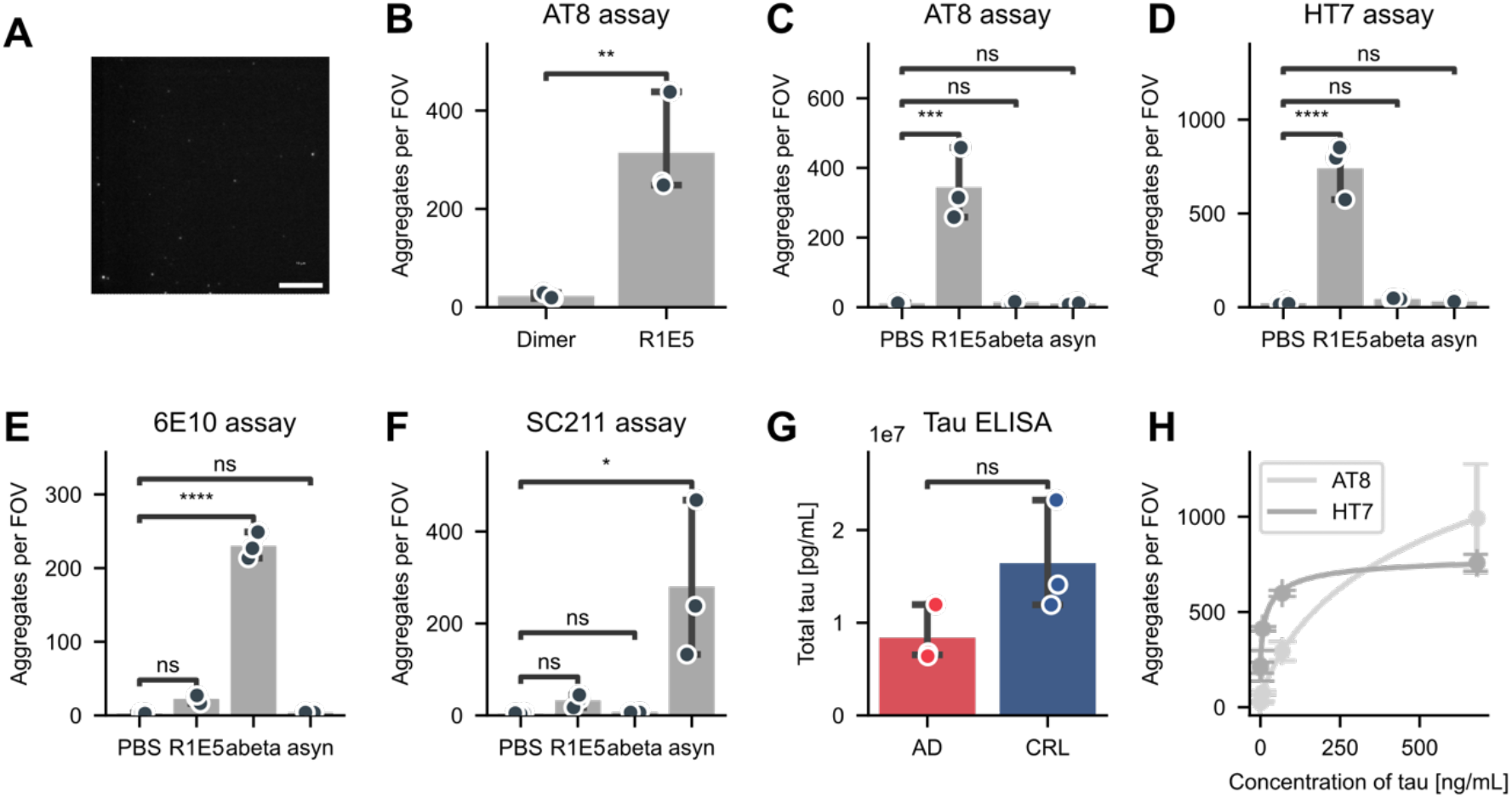
Validation of the tau SiMPull assay. **(A)** Representative image of a sample containing no tau (BSA) where no fluorescent spots are quantified. **(B)** Quantification of fluorescent spots observed using the HT7-epitope dimer-mimicking peptide in an AT8 tau SiMPull assay. As a positive control, a cell lysate from HEK cells overexpressing hyperphosphorylated tau aggregates (R1E5) was used. **(C-D)** SiMPull assays against **(C)** AT8 or **(D)** HT7 were tested for cross-reactivity with recombinant amyloid-β and α-synuclein aggregates. **(E-F)** SiMPull assays against **(E)** amyloid-β and **(F)** α-synuclein demonstrate recombinant aggregates of amyloid-β and α-synuclein can be successfully detected by their respective SiMPull assays. **(G)** Total tau concentration in AD and control brain homogenate samples determined using a commercial tau ELISA kit (not aggregate specific). **(H)** Four-parameter logistic regression fitted to the number of aggregates per FOV for a serial dilution of a highly concentrated tau aggregate sample (Braak VI brain homogenate). Panel B - H show the mean ± S.D. of n=3 technical replicates compared using a t-Test (panel B, G) or a one-way Anova with post-hoc Tukey HSD test (panel C-F). ns: *p* > 0.05, *: *p* < 0.05, **: *p* < 0.01, ***: *p* < 0.001, ****: *p* < 0.0001.

**Supplementary Fig. 2.**
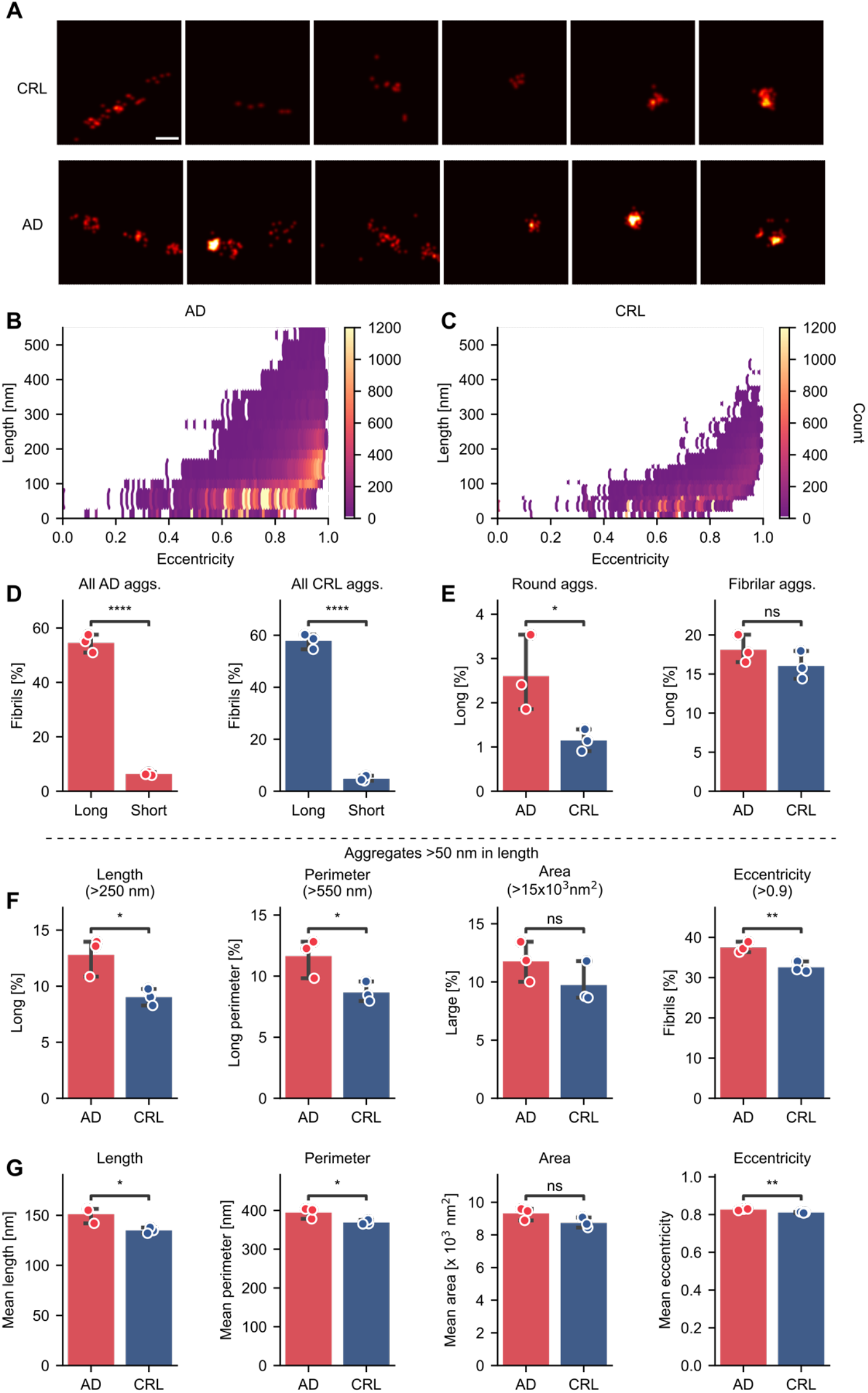
Morphological characterization of super-resolved tau aggregates derived from human brain homogenate. **(A)** Representative images of aggregates observed in human brain homogenate from AD and control patients, showing fibrillar aggregates and smaller, round aggregates. Scale bar = 100 nm. **(B)** Length and eccentricity of the individual aggregates in AD brain homogenate samples. **(C)** Length and eccentricity of the individual aggregates in control brain homogenate samples. **(D)** Percentage of fibrils (eccentricity >0.9) in aggregates classified as either long (>250 nm) or short (<100 nm) in AD (red) and control brain (blue) homogenate. **(E)** Percentage of long aggregates (>250 nm) in aggregates classified as round (eccentricity <0.7) or fibrillar (eccentricity >0.9) aggregates in AD (red) and CRL (blue) brain homogenate. **(F)** Percentage of large and fibrillar aggregates (length: >250 nm, perimeter: > 550 nm, area > 15x 10^3^ nm^2^, eccentricity > 0.9) after filtering out aggregates <50 nm in length. **(G)** Mean length, perimeter, area and eccentricity of aggregates after filtering out aggregates <50 nm in length. Panel D - I show the mean ± S.D. of n=3 biological replicates compared using a t-Test. ns: *p* > 0.05, *: *p* < 0.05, **: *p* < 0.01, ****: *p* < 0.0001.

**Supplementary Fig. 3.**
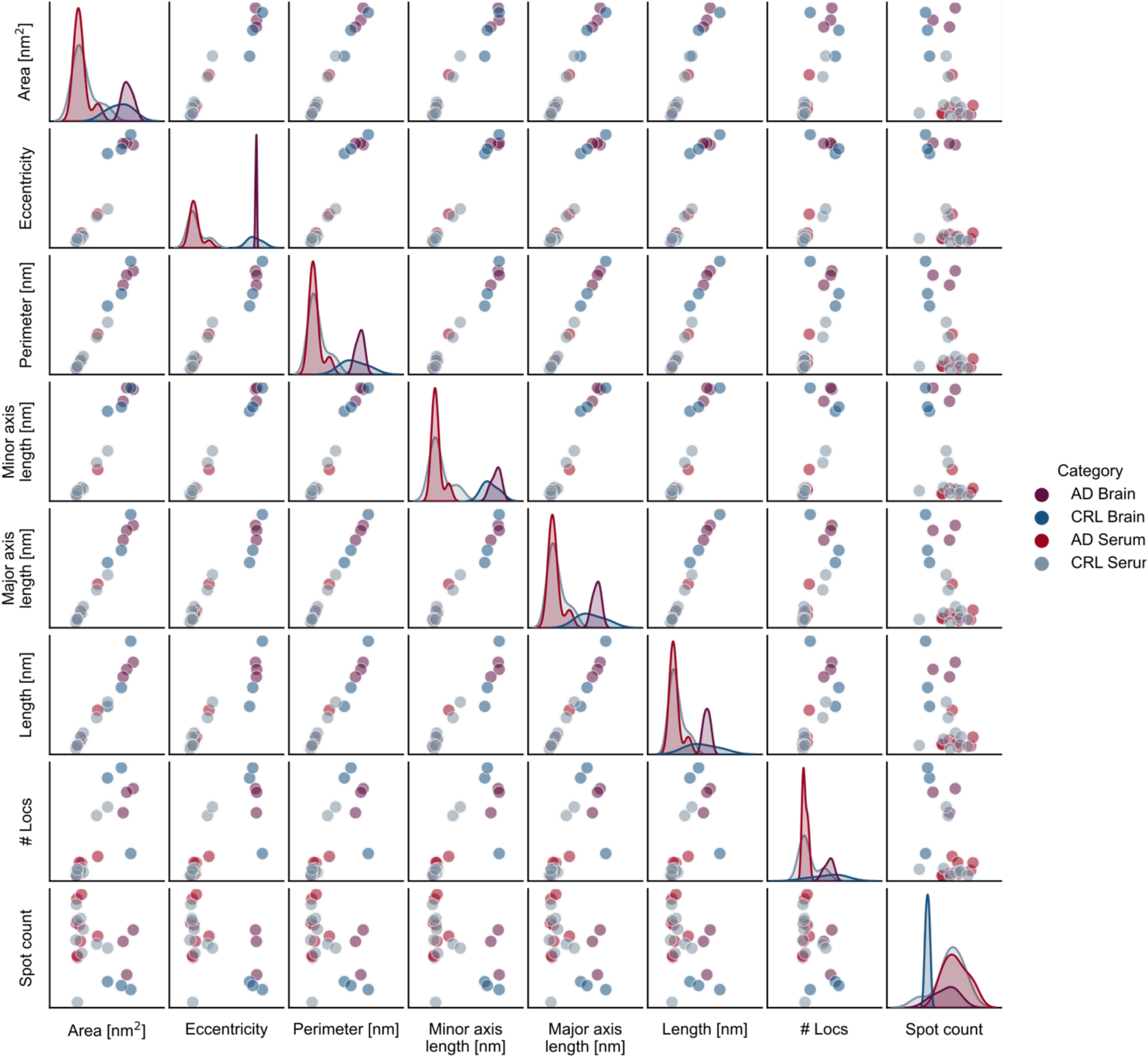
Pairwise comparison of morphological features of tau aggregates from brain homogenate and serum. Pairwise relationship of morphological features of tau aggregates (HT7) in human brain homogenate and serum.

**Supplementary Fig. 4.**
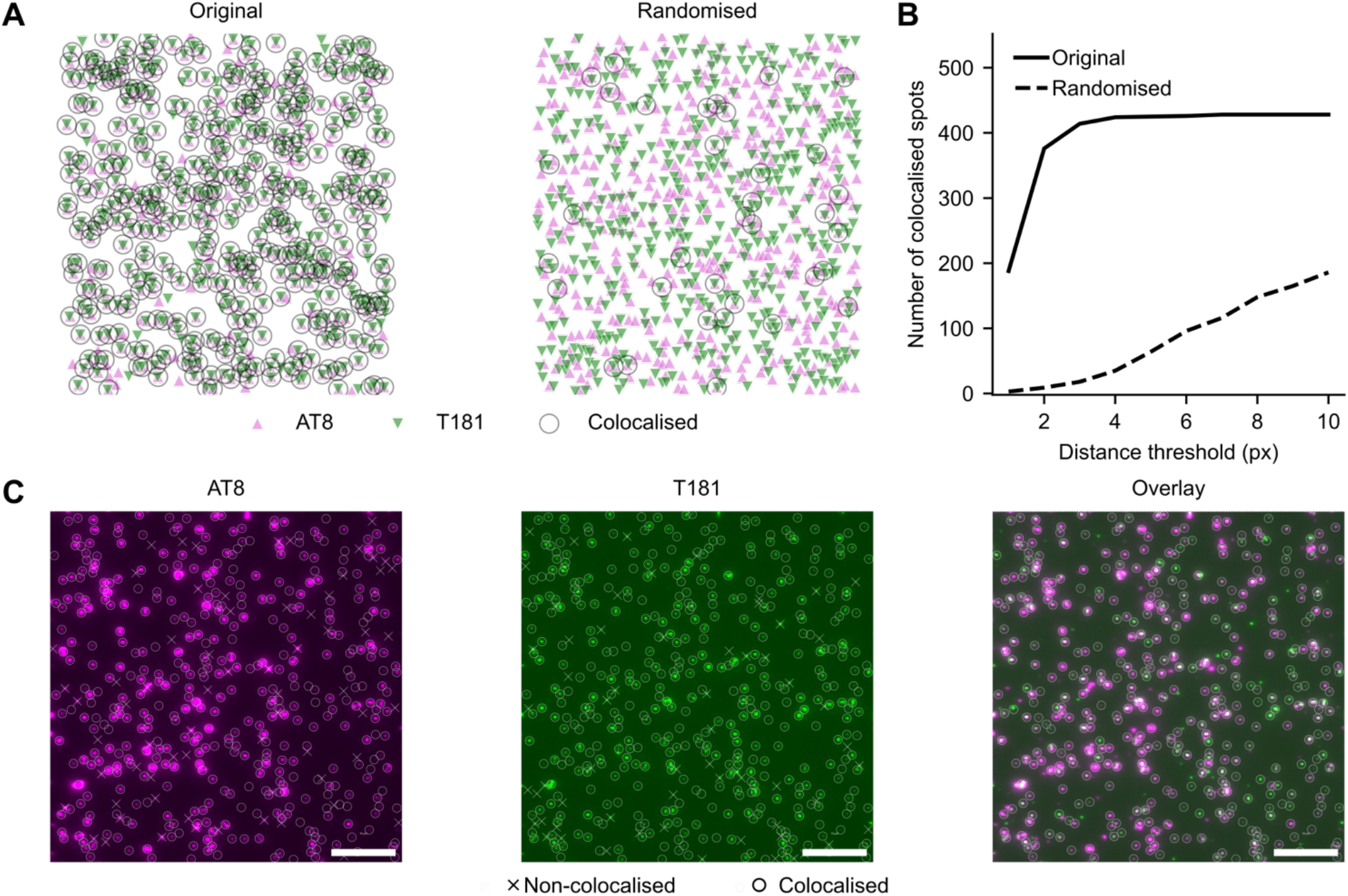
Co-localization analysis. **(A)** Exemplar visualization of spots detected by ComDet in each channel (in this case, AF647-labelled AT8 △ and AF488-labelled T181 ∇). Spots for a given channel are compared with those from the opposing channel, and pairs of spots for which the Euclidean distance is less than the threshold value are considered co-localized (O). In the event that more than one spot passes the threshold, the spot with the shortest distance is selected. To estimate the likelihood of these spots being co-localized by chance, the second channel coordinates are inverted, and the co-localization calculation repeated (Randomized). **(B)** The number of co-localized spots as calculated for the original or randomized spots shown in A at threshold distances ranging from 1 to 10. **(C)** Detected spots shown in A overlayed onto the source AT8 or T181 images, demonstrating those which were found to be co-localized (O) or non-colocalized (X). Scale bar = 10 μm.

**Supplementary Fig. 5.**
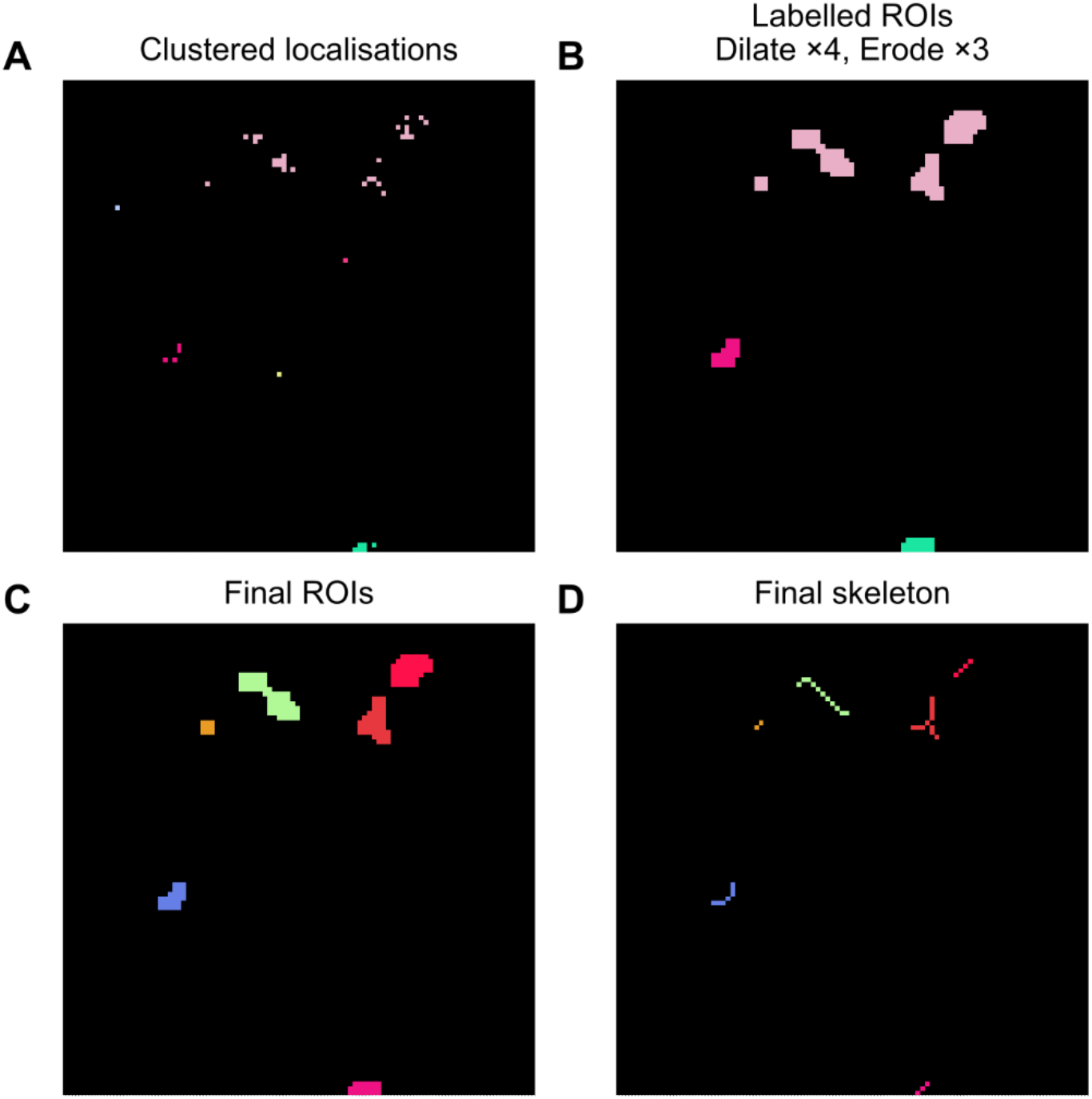
Super-resolution measurement of individual molecules. **(A)** Localizations are first clustered using DBSCAN with permissive parameters to discard isolated localizations (∼noise). **(B)** The resultant localizations are subjected to rounds of morphological dilation, closing and erosion to arrive at single connected regions of interest (ROIs). **(C)** The resultant ROIs are then relabeled such that individual ROIs are given a single unique identifier. **(D)** Each ROI is skeletonized to allow for length measurements. In all panels, pixel color represents an arbitrary pixel/object identifier.

**Table 1.** Brain Homogenate Patient Information.

| **Subject ID** | **Sex** | **Brain region** | **Group** | **Age at**<br>**Diagnosis**<br>**[years]** | **Age at**<br>**Death**<br>**[years]** | **Braak**<br>**Stage** |
| --- | --- | --- | --- | --- | --- | --- |
| NP16.00028 | M | Frontal cortex | Control |  | 68 | I |
| NP18.00159 | F | Frontal cortex | Control |  | 68 | 0 |
| NP19.00009 | M | Frontal cortex | Control |  | 66 | I |
| NP14.00055 | F | Frontal cortex | AD | 64 | 70 | VI |
| NP17.00246 | F | Frontal cortex | AD | 67 | 74 | VI |
| NP18.00042 | F | Frontal cortex | AD | 86 | 100 | VI |

**Table 2.** Serum Patient Information (CRL = control, AD = Alzheimer’s disease).

| **Group** | **Sex** | **Age at visit**<br>**[years]** | **Symptom**<br>**duration**<br>**[years]** | **Diagnosis**<br>**duration**<br>**[years]** |
| --- | --- | --- | --- | --- |
| CRL1 | M | 81.0 |  |  |
| CRL2 | F | 70.0 |  |  |
| CRL3 | F | 65.0 |  |  |
| CRL4 | F | 56.0 |  |  |
| CRL5 | F | 73.0 |  |  |
| CRL6 | M | 67.0 |  |  |
| CRL7 | M | 81.0 |  |  |
| CRL8 | M | 87.0 |  |  |
| CRL9 | M | 54.0 |  |  |
| CRL10 | M | 88.0 |  |  |
| AD1 | F | 60.3 | 2.8 | 2.3 |
| AD2 | F | 62.3 | 5.9 | 2.3 |
| AD3 | F | 53.8 | 3.3 | 1.7 |
| AD4 | F | 59.8 | 5.7 | 3.7 |
| AD5 | M | 62.3 | 3.7 | 2.1 |
| AD6 | F | 68.9 | 8.1 | 1.6 |
| AD7 | F | 78.1 | 2.5 | 0.7 |
| AD8 | M | 56.5 | 7.1 | 1.1 |
| AD9 | M | 59.6 | 7.1 | 2.1 |
| AD10 | F | 57.5 | 6.6 | 2.1 |

## References

1. Flament, S., Delacourte, A., Verny, M., Hauw, J. J. & Javoy-Agid, F. Abnormal Tau proteins in progressive supranuclear palsy. Similarities and differences with the neurofibrillary degeneration of the Alzheimer type. Acta Neuropathol 81, 591–596 (1991).

2. Dickson, D. W. Pick’s Disease: A Modern Approach. Brain Pathology 8, 339 (1998).

3. Goedert, M. Tau protein and the neurofibrillary pathology of Alzheimer’s disease. Trends Neurosci 16, 460–465 (1993).

4. Grundke-Iqbal, I. et al. Abnormal phosphorylation of the microtubule-associated protein tau (tau) in Alzheimer cytoskeletal pathology. Proceedings of the National Academy of Sciences 83, 4913–4917 (1986).

5. Drubin, D. G. & Kirschner, M. W. Tau protein function in living cells. J Cell Biol 103, 2739–2746 (1986).

6. Weingarten, M. D., Lockwood, A. H., Hwo, S. Y. & Kirschner, M. W. A protein factor essential for microtubule assembly. Proc Natl Acad Sci U S A 72, 1858–1862 (1975).

7. Iqbal, K., Liu, F., Gong, C.-X. & Grundke-Iqbal, I. Tau in Alzheimer Disease and Related Tauopathies. Curr Alzheimer Res 7, 656 (2010).

8. Kametani, F. & Hasegawa, M. Reconsideration of Amyloid Hypothesis and Tau Hypothesis in Alzheimer’s Disease. Front Neurosci 12, (2018).

9. Bejanin, A. et al. Tau pathology and neurodegeneration contribute to cognitive impairment in Alzheimer’s disease. Brain 140, 3286–3300 (2017).

10. Arriagada, P. V., Growdon, J. H., Hedley-Whyte, E. T. & Hyman, B. T. Neurofibrillary tangles but not senile plaques parallel duration and severity of Alzheimer’s disease. Neurology 42, 631–639 (1992).

11. Kopeikina, K. J., Hyman, B. J. & Spires-Jones, T. L. Soluble forms of tau are toxic in Alzheimer’s disease. Transl Neurosci 3, 223 (2012).

12. Lasagna-Reeves, C. A. et al. Tau oligomers impair memory and induce synaptic and mitochondrial dysfunction in wild-type mice. Mol Neurodegener 6, (2011).

13. Lasagna-Reeves, C. A. et al. Alzheimer brain-derived tau oligomers propagate pathology from endogenous tau. Scientific Reports 2012 2:1 2, 1–7 (2012).

14. Jucker, M. & Walker, L. C. Propagation and spread of pathogenic protein assemblies in neurodegenerative diseases. Nat Neurosci 21, 1341–1349 (2018).

15. Fitzpatrick, A. W. P. et al. Cryo-EM structures of tau filaments from Alzheimer’s disease. Nature 2017 547:7662 547, 185–190 (2017).

16. Shi, Y. et al. Structure-based classification of tauopathies. Nature 2021 598:7880 598, 359–363 (2021).

17. Hitt, B. D. et al. Ultrasensitive tau biosensor cells detect no seeding in Alzheimer’s disease CSF. Acta Neuropathol Commun 9, 1–10 (2021).

18. Saijo, E. et al. Ultrasensitive and selective detection of 3-repeat tau seeding activity in Pick disease brain and cerebrospinal fluid. Acta Neuropathol 133, 751–765 (2017).

19. Kraus, A. et al. Seeding selectivity and ultrasensitive detection of tau aggregate conformers of Alzheimer disease. Acta Neuropathol 137, 585–598 (2019).

20. Maeda, S. et al. Granular tau oligomers as intermediates of tau filaments. Biochemistry 46, 3856–3861 (2007).

21. Jain, A., Liu, R., Xiang, Y. K. & Ha, T. Single-molecule pull-down for studying protein interactions. Nat Protoc 7, (2012).

22. Arbaciauskaite, M., Lei, Y. & Cho, Y. K. High-specificity antibodies and detection methods for quantifying phosphorylated tau from clinical samples. Antib Ther 4, 34 (2021).

23. Whiten, D. R. et al. Nanoscopic Characterisation of Individual Endogenous Protein Aggregates in Human Neuronal Cells. ChemBioChem 19, 2033–2038 (2018).

24. Rust, M. J., Bates, M. & Zhuang, X. Sub-diffraction-limit imaging by stochastic optical reconstruction microscopy (STORM). Nature Methods 2006 3:10 3, 793–796 (2006).

25. Dugger, B. N. et al. The Presence of Select Tau Species in Human Peripheral Tissues and Their Relation to Alzheimer’s Disease. J Alzheimers Dis 51, 345–356 (2016).

26. Barthélemy, N. R., Horie, K., Sato, C. & Bateman, R. J. Blood plasma phosphorylated-tau isoforms track CNS change in Alzheimer’s disease. J Exp Med 217, (2020).

27. Barthélemy, N. R. et al. Cerebrospinal fluid phospho-tau T217 outperforms T181 as a biomarker for the differential diagnosis of Alzheimer’s disease and PET amyloid-positive patient identification. Alzheimers Res Ther 12, (2020).

28. Jicha, G. A., Bowser, R., Kazam, I. G. & Davies, P. Alz-50 and MC-1, a new monoclonal antibody raised to paired helical filaments, recognize conformational epitopes on recombinant tau. J Neurosci Res 48, 128–132 (1997).

29. de Silva, R. et al. An immunohistochemical study of cases of sporadic and inherited frontotemporal lobar degeneration using 3R- and 4R-specific tau monoclonal antibodies. Acta Neuropathol 111, 329–340 (2006).

30. Suárez-Calvet, M. et al. Novel tau biomarkers phosphorylated at T181, T217 or T231 rise in the initial stages of the preclinical Alzheimer’s continuum when only subtle changes in Aβ pathology are detected. EMBO Mol Med 12, (2020).

31. Qin, W. et al. Phosphorylated Tau 181 Serum Levels Predict Alzheimer’s Disease in the Preclinical Stage. Front Aging Neurosci 14, 559 (2022).

32. Wesseling, H. et al. Tau PTM Profiles Identify Patient Heterogeneity and Stages of Alzheimer’s Disease. Cell 183, 1699–1713.e13 (2020).

33. Dujardin, S. et al. Tau molecular diversity contributes to clinical heterogeneity in Alzheimer’s disease. Nature Medicine 2020 26:8 26, 1256–1263 (2020).

34. Neddens, J. et al. Phosphorylation of different tau sites during progression of Alzheimer’s disease. Acta Neuropathol Commun 6, 52 (2018).

35. Blömeke, L. et al. Quantitative detection of α-Synuclein and Tau oligomers and other aggregates by digital single particle counting. npj Parkinson’s Disease 2022 8:1 8, 1–13 (2022).

36. Demmerle, J., Wegel, E., Schermelleh, L. & Dobbie, I. M. Assessing resolution in super-resolution imaging. Methods 88, 3–10 (2015).

37. Chen, X., Chen, M., Schafer, N. P. & Wolynes, P. G. Exploring the interplay between fibrillization and amorphous aggregation channels on the energy landscapes of tau repeat isoforms. Proc Natl Acad Sci U S A 117, 4125–4130 (2020).

38. Meng, J. X. et al. Hyperphosphorylated tau self-assembles into amorphous aggregates eliciting TLR4-dependent responses. Nat Commun 13, (2022).

39. Corbett, G. T. et al. PrP is a central player in toxicity mediated by soluble aggregates of neurodegeneration-causing proteins. Acta Neuropathol 139, 503–526 (2020).

40. Cremades, N. et al. Direct observation of the interconversion of normal and toxic forms of α-synuclein. Cell 149, 1048–1059 (2012).

41. Goedert, M., Spillantini, M. G., Cairns, N. J. & Crowther, R. A. Tau proteins of Alzheimer paired helical filaments: abnormal phosphorylation of all six brain isoforms. Neuron 8, 159–168 (1992).

42. Chandradoss, S. D. et al. Surface passivation for single-molecule protein studies. J Vis Exp (2014) doi:10.3791/50549.

43. Jain, A., Liu, R., Xiang, Y. K. & Ha, T. Single-molecule pull-down for studying protein interactions. Nat Protoc 7, 445–452 (2012).

44. Edelstein, A. D. et al. Advanced methods of microscope control using μManager software. J Biol Methods 1, e10 (2014).

45. Katrukha, E. ekatrukha/ComDet: ComDet 0.5.3. (2020) doi:10.5281/ZENODO.4281064.

46. Schnitzbauer, J., Strauss, M. T., Schlichthaerle, T., Schueder, F. & Jungmann, R. Super-resolution microscopy with DNA-PAINT. Nature Protocols 2017 12:6 12, 1198–1228 (2017).

47. Pedregosa FABIANPEDREGOSA, F. et al. Scikit-learn: Machine Learning in Python. The Journal of Machine Learning Research 12, 2825–2830 (2011).

48. Van Der Walt, S. et al. scikit-image: image processing in Python. PeerJ 2, (2014).

49. Nunez-Iglesias, J., Blanch, A. J., Looker, O., Dixon, M. W. & Tilley, L. A new Python library to analyse skeleton images confirms malaria parasite remodelling of the red blood cell membrane skeleton. PeerJ 2018, e4312 (2018).

50. Virtanen, P. et al. SciPy 1.0: fundamental algorithms for scientific computing in Python. Nature Methods 2020 17:3 17, 261–272 (2020).

51. Charlier, F. et al. trevismd/statannotations: v0.5. (2022) doi:10.5281/ZENODO.7213391.

